# Implications for Faster Healing with Fewer Applications: Sustained Graft Persistence and Favorable Wound-Healing Biology of a Proprietary Processed Placental Tissue Allograft

**DOI:** 10.64898/2026.10.14.759179

**Authors:** Taylor Carson, Joydeep Basu

**Affiliations:** Applied Biologics LLC, 8125 N 86^th^ Pl., Scottsdale, AZ 85258

## Abstract

**Introduction:** Placental tissue allografts are widely utilized as wound coverings and barriers. In addition to tissue source, manufacturing and processing methodologies can influence extracellular matrix (ECM) architecture, mechanical stability, protease susceptibility, degradation kinetics, and graft persistence following implantation. An optimized placental tissue allograft should maintain sufficient matrix integrity during the active period of tissue repair while preserving the biological characteristics associated with a favorable wound-healing environment. Our previously published work demonstrated that a proprietary processed human amniotic membrane allograft exhibited a more favorable wound-healing biologic profile than conventionally processed placental tissue. The finding that no XWRAP-treated wound had reached the reapplication threshold at Day 14, considered together with previously demonstrated favorable wound-healing biology, provides a mechanistic rationale for the hypothesis that XWRAP may support faster healing with fewer applications over a course of wound management.

**Methods:** Graft integrity was evaluated every 2 days for 14 days following application to standardized full-thickness excisional wounds in Sprague-Dawley rats. XWRAP, a proprietary processed human amniotic membrane allograft, was compared with a donor-matched conventionally processed human amniotic membrane comparator and a commercial placental tissue comparator. Graft integrity was scored from 0 to 5, where 0 represented complete graft loss or resorption and 5 represented an intact graft. A score ≤1 represented the protocol-defined threshold for reapplication during non-terminal visits. Longitudinal graft-integrity trajectories, original-graft degradation, documented reapplications, and the proportion of wounds reaching the reapplication threshold were evaluated. At each visit, graft integrity in XWRAP-treated wounds was compared with the pooled comparator groups (primary comparison) and with each comparator separately using animal-level Mann–Whitney tests, with Holm adjustment across visits; reapplication frequency was compared using exact Poisson rate ratios.

**Results:** XWRAP demonstrated markedly greater late-phase persistence than both comparator products. At Day 14, mean graft-integrity score was 3.88 for XWRAP compared with 0.93 for donor-matched conventionally processed amnion and 1.69 for the commercial comparator. From Days 10 through 14, XWRAP graft integrity was significantly greater than both comparators (all p≤0.001), with Cliff’s δ values of 0.95–1.00. Compared with the pooled comparator groups, XWRAP graft integrity was significantly greater from Day 8 (3.31 vs. 2.93; p=0.022) through Day 14 (3.88 vs. 1.33; p<0.001); differences at Days 10–14 remained significant after Holm adjustment, and no significant differences were observed during Days 2–6.

At the terminal Day 14 assessment, none of the XWRAP-treated wounds had reached the protocol-defined reapplication threshold (0/16; 0%), compared with 11/14 (79%) wounds treated with donor-matched conventional amnion and 5/16 (31%) wounds treated with the commercial comparator. Analysis limited to original grafts demonstrated the same persistence pattern, with Day 14 integrity scores of 3.92, 0.90, and 1.83, respectively. The proportion of wounds at the reapplication threshold at Day 14 was significantly lower with XWRAP than with the pooled comparators (0/16 vs. 16/30; p<0.001). Documented reapplications, all of which occurred during Days 2–6, did not differ between XWRAP and the pooled comparators (3.24 vs. 3.40 per 100 wound-days; rate ratio 0.95, 95% CI 0.33–2.53; p=1.00).

**Discussion:** Proprietary processing produced a substantially different graft-persistence profile, with XWRAP remaining largely intact while comparator tissues progressed toward resorption. Importantly, this greater graft persistence complements our previously published findings demonstrating that XWRAP also produced a more favorable wound-healing biologic profile across multiple measures of cellular proliferation, inflammation, tissue injury, and wound resolution. Taken together, these findings support two complementary attributes of XWRAP: sustained graft persistence and favorable wound-healing biology. The finding that no XWRAP-treated wound had reached the reapplication threshold at Day 14 provides a mechanistic rationale for the hypothesis that this combination may extend application intervals and reduce application requirements over a course of wound management.

## Introduction

Cutaneous wound repair is a highly coordinated biological process involving inflammatory cells, resident stromal cells, extracellular matrix, soluble signaling molecules, and mechanical cues. Successful repair requires progression through overlapping phases of hemostasis, inflammation, proliferation, and remodeling. During this process, the extracellular matrix provides structural support and a dynamic biological environment for cellular migration, proliferation, angiogenesis, matrix deposition, and progressive tissue remodeling. [2–7]

Human amniotic membrane has been utilized clinically for more than a century. [8] Its structural properties arise from a complex extracellular matrix containing multiple collagen isoforms, laminin, fibronectin, hyaluronic acid, proteoglycans, and other matrix components. Amnion also contains matrix-associated cytokines and growth factors that have been investigated for their potential roles in cellular adhesion, migration, proliferation, angiogenesis, inflammatory regulation, and tissue remodeling. [9–17]

In addition to these biological characteristics, human amniotic membrane provides a physical covering and barrier. The duration for which the matrix remains intact following application may therefore represent an important product characteristic. A graft that undergoes rapid degradation, resorption, displacement, or removal may require replacement to maintain tissue coverage. Conversely, a graft that remains intact through a greater portion of the wound-management interval may reduce the frequency with which additional tissue must be applied.

Native human amnion is mechanically fragile and biologically labile. Numerous preservation and manufacturing methods have therefore been developed to modify tissue handling, storage, mechanical stability, and resistance to degradation. These include cryopreservation, dehydration, lyophilization, decellularization, terminal sterilization, supercritical carbon dioxide processing, and chemical modification of extracellular matrix components. [18–20]

Chemical modification has received particular attention because the mechanical properties and degradation kinetics of collagen-rich matrices can be altered through intermolecular stabilization of the ECM. Crosslinking approaches described in the scientific literature include glutaraldehyde, carbodiimide-based chemistries, genipin, and naturally occurring polyphenols. Depending upon the chemistry and degree of modification, these approaches can increase collagen stability, mechanical strength, and resistance to enzymatic degradation. [21–30]

The relationship between stabilization and biological performance, however, is not simply linear. Insufficient stabilization may permit premature degradation within a protease-rich environment, shortening the period during which the matrix remains structurally present. Excessive stabilization may reduce matrix porosity, alter cellular attachment sites, impede host-cell infiltration, modify release of matrix-associated signaling molecules, or interfere with progressive host remodeling. Accordingly, the objective of optimized matrix processing is not maximal persistence, but an appropriate balance between matrix stability, biologic activity, and progressive remodeling. [26–30]

Manufacturing methodology may therefore influence two complementary dimensions of placental tissue performance: what the matrix does biologically while present and how long the matrix remains present at the tissue interface.

We previously evaluated XWRAP, a proprietary processed human amniotic membrane allograft, in a standardized rodent full-thickness excisional wound model [1]. Compared with conventionally processed placental tissue, XWRAP demonstrated a more favorable wound-healing biologic profile, including increased proliferative activity, reduced ulceration severity, reduced giant-cell response, favorable inflammatory remodeling, and more advanced progression toward wound resolution [1].

The present analysis addresses the complementary question of graft persistence and application utilization. Using longitudinal observations from the same experimental model, we evaluated graft-integrity trajectories, degradation kinetics, documented reapplications, and the proportion of wounds reaching a predefined reapplication threshold.

We hypothesized that proprietary processing would produce greater late-phase graft persistence than conventionally processed placental tissue. When considered together with the previously reported wound-healing findings, greater persistence could provide a mechanistic basis for maintaining favorable wound-healing biology over a longer interval and potentially reducing the number of applications required during a course of wound management.

## Materials and Methods

### Preclinical Animal Model

Rodent studies were conducted using male Sprague-Dawley rats in a standardized full-thickness excisional skin wound model. [31] Each animal received two dorsal full-thickness excisional wounds, designated cranial and caudal. Test articles were applied directly to the wound bed and secured with gauze, bandaging, and protective wrapping. The present analysis focused on three treatment groups from an original 5 arm trial-Group 5 (G5): XWRAP, proprietary processed human amniotic membrane allograft; Group 3 (G3): conventionally processed human amniotic membrane comparator derived from the same donor source as XWRAP; and Group 4 (G4): commercial placental tissue comparator. G5 and G3 were derived from the same amnion donor (00597), permitting comparison of differently processed tissue originating from a common donor source. G4 represented an independent commercial comparator. Each treatment group initially contained 10 animals and 20 wounds. At Day 14, 16 wounds remained evaluable in G5, 14 in G3, and 16 in G4.

### Longitudinal Graft-Integrity Assessment

Wounds were evaluated every 2 days from Day 0 through Day 14. At each visit, residual graft integrity was scored using an ordinal scale from 0 to 5, where 5 represented an intact graft and 0 represented complete graft loss or resorption. A graft-integrity score ≤1 represented the protocol-defined threshold for reapplication during Days 2–12. Day 14 represented the terminal study assessment. Wounds reaching the reapplication threshold at Day 14 therefore did not receive an additional graft.

### Reapplication Analysis

Individual wound records were reviewed at each study visit for placement of a new graft. A reapplication event was defined as placement of an additional graft during Days 2–12. From a mechanistic standpoint, graft loss associated with in-situ degradation or resorption is distinct from mechanical loss associated with dressing adhesion or displacement. From an application-utilization standpoint, however, either mechanism may result in loss of adequate graft coverage and the practical requirement for another application. Documented reapplications were therefore retained irrespective of the recorded mechanism of graft loss.

To separately characterize persistence of the original graft, an additional analysis censored each wound after its first reapplication. This permitted evaluation of original-graft degradation independently of subsequent replacement material.

### Statistical Analysis

Because each animal contributed cranial and caudal wounds, paired wound scores were averaged for primary animal-level comparisons. Per-time-point comparisons were performed using two-sided Mann–Whitney U tests. Mean treatment differences were calculated with 95% bootstrap confidence intervals.

Effect size was characterized using Cliff’s δ. Absolute values ≥0.147 were considered small, ≥0.33 medium, and ≥0.474 large. Sensitivity analyses included wound-level comparisons and longitudinal models incorporating study day and treatment with cluster-robust standard errors by animal. Original-graft degradation was additionally evaluated using longitudinal degradation slopes. Holm adjustment was used to evaluate robustness across repeated time-point comparisons.

The primary comparison was XWRAP (G5) versus the pooled comparator groups (G3 and G4 combined). XWRAP was also compared separately with donor-matched conventional amnion (G3) and with the commercial comparator (G4), and the two comparator groups were compared with each other for context. A two-sided p<0.05 was considered statistically significant, and 0.05≤p<0.10 was reported as a trend toward significance. Holm adjustment was applied across the seven post-baseline visits within each comparison.

Mean graft integrity over predefined intervals (Days 2–6, 8–10, 10–14 and 2–14) and within-wound change in integrity between Day 6 and Day 14 were compared at the animal level using Mann–Whitney tests. Original-graft degradation slopes were estimated for each wound followed to Day 14 by least-squares regression of integrity score on study day and averaged per animal. Time to first original-graft reapplication (integrity score ≤1) was estimated using the Kaplan–Meier method and compared using the log-rank test. Reapplication frequency was expressed per 100 wound-days at risk (Day 0 to the last assessment, censored at Day 12) and compared using exact Poisson rate ratios, with a supporting animal-level cluster-bootstrap confidence interval (10,000 resamples). Per-visit reapplication proportions and the proportion of wounds at or below the reapplication threshold at Day 14 were compared using Fisher’s exact test. Analyses were performed in Python (pandas and SciPy).

## Results

### Longitudinal Graft-Persistence Trajectories

All grafts were intact at application, with an integrity score of 5.0 at Day 0. During the early study period, graft-integrity scores decreased across treatment groups. Through Day 6, no sustained persistence advantage was evident for XWRAP. Beginning at Day 8, however, the treatment trajectories separated. Mean XWRAP integrity was 3.31 at Day 8, compared with 3.00 for donor-matched conventional amnion and 2.88 for the commercial comparator.

During this early period, mean XWRAP integrity was numerically lower than that of the pooled comparator groups at Day 2 (2.70 vs. 3.40; p=0.141) and Day 6 (2.85 vs. 3.26; p=0.155) and similar at Day 4 (3.25 vs. 2.97; p=1.000). At Day 6, XWRAP integrity was significantly lower than that of the commercial comparator (2.85 vs. 3.55; p=0.025) but did not differ from donor-matched conventional amnion (2.94; p=0.964). Group trajectories are shown in Figure 1 and Table 1, and individual wound trajectories in Figure 2.

**Figure 1.**
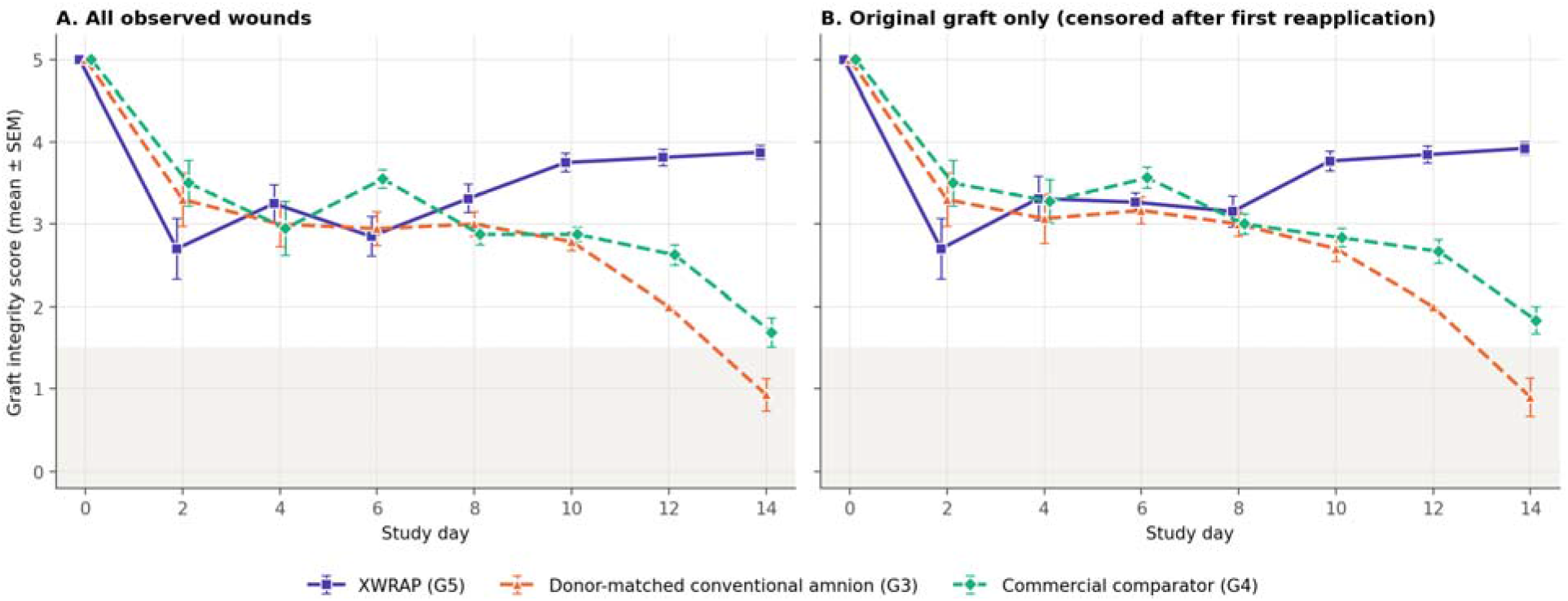
Longitudinal graft-integrity trajectories by treatment group (mean ± SEM of wound scores). (A) All observed wounds. (B) Original graft only, with each wound censored after its first reapplication. The shaded band indicates the protocol-defined reapplication threshold (score ≤1).

**Figure 2.**
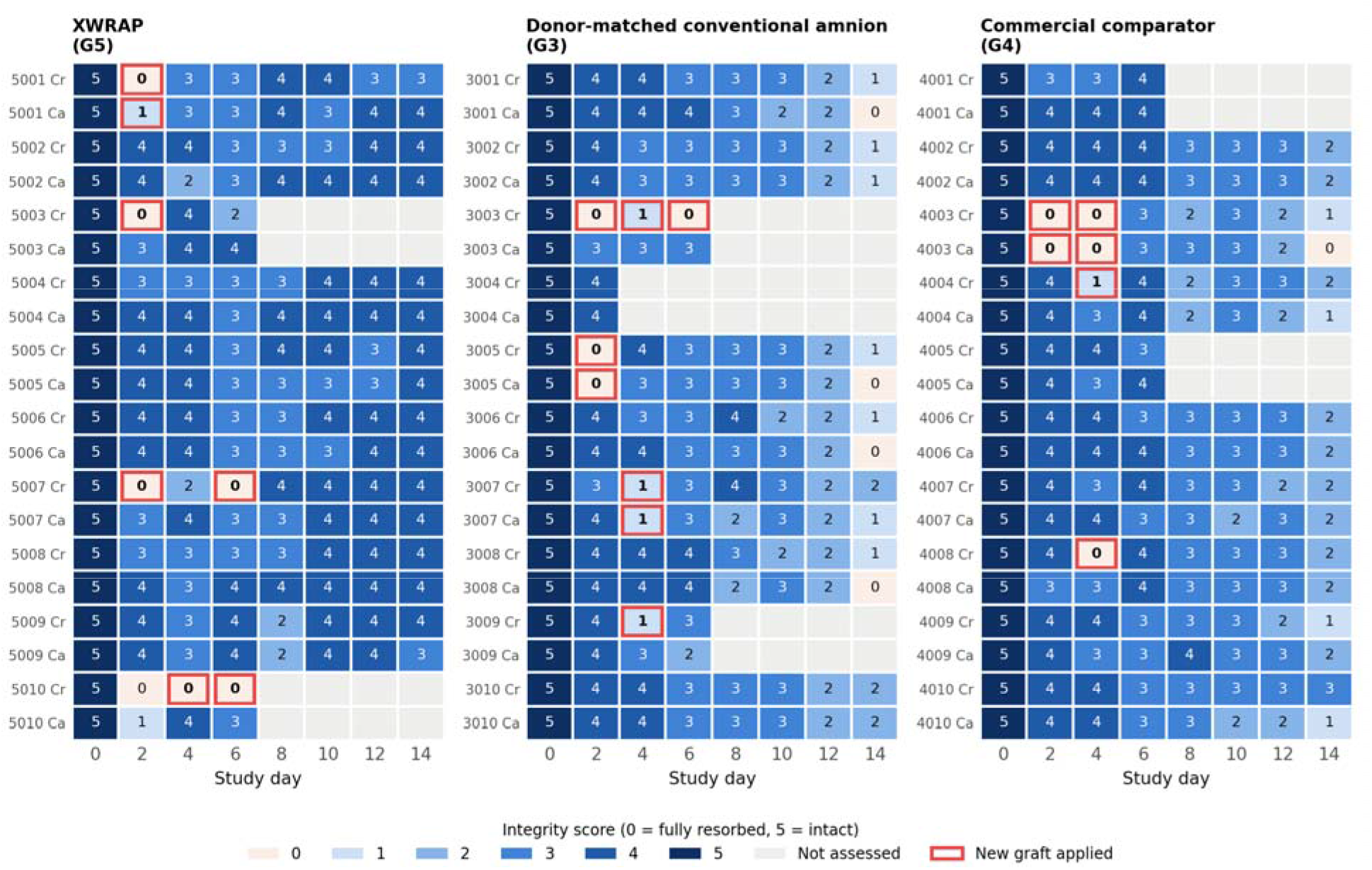
Individual wound graft-integrity scores by study day for each treatment group (Cr, cranial; Ca, caudal). Red outlines indicate visits at which a new graft was applied; grey cells indicate wounds not assessed.

**Table 1.** Graft-integrity score by treatment group and study day, all observed wounds: mean ± SD (number of wounds).

| Group | Day 0 | Day 2 | Day 4 | Day 6 | Day 8 | Day 10 | Day 12 | Day 14 |
| --- | --- | --- | --- | --- | --- | --- | --- | --- |
| XWRAP (G5) | 5.00 ± 0.00 (20) | 2.70 ± 1.66 (20) | 3.25 ± 1.02 (20) | 2.85 ± 1.09 (20) | 3.31 ± 0.70 (16) | 3.75 ± 0.45 (16) | 3.81 ± 0.40 (16) | 3.88 ± 0.34 (16) |
| Donor-matched conventional amnion (G3) | 5.00 ± 0.00 (20) | 3.30 ± 1.45 (20) | 3.00 ± 1.19 (18) | 2.94 ± 0.87 (18) | 3.00 ± 0.55 (14) | 2.79 ± 0.43 (14) | 2.00 ± 0.00 (14) | 0.93 ± 0.73 (14) |
| Commercial comparator (G4) | 5.00 ± 0.00 (20) | 3.50 ± 1.24 (20) | 2.95 ± 1.47 (20) | 3.55 ± 0.51 (20) | 2.88 ± 0.50 (16) | 2.88 ± 0.34 (16) | 2.63 ± 0.50 (16) | 1.69 ± 0.70 (16) |
| Pooled comparators (G3+G4) | 5.00 ± 0.00 (40) | 3.40 ± 1.34 (40) | 2.97 ± 1.33 (38) | 3.26 ± 0.76 (38) | 2.93 ± 0.52 (30) | 2.83 ± 0.38 (30) | 2.33 ± 0.48 (30) | 1.33 ± 0.80 (30) |

At Day 10, XWRAP remained at 3.75, whereas donor-matched conventional amnion and the commercial comparator were 2.79 and 2.88, respectively. At Day 12, XWRAP remained at 3.81, compared with 2.00 for conventional amnion and 2.63 for the commercial comparator. By Day 14, the difference was pronounced. XWRAP maintained a mean integrity score of 3.88, while donor-matched conventional amnion had declined to 0.93 and the commercial comparator to 1.69. Corresponding means for the pooled comparator groups were 2.93 at Day 8, 2.83 at Day 10, 2.33 at Day 12, and 1.33 at Day 14 (Table 1).

### XWRAP Demonstrated Substantially Greater Late-Phase Graft Integrity

In the primary comparison with the pooled comparator groups, XWRAP graft integrity was significantly greater at Day 8 (3.31 vs. 2.93; p=0.022; Cliff’s δ=0.56), Day 10 (3.75 vs. 2.83; p<0.001), Day 12 (3.81 vs. 2.33; p<0.001), and Day 14 (3.88 vs. 1.33; p<0.001), with Cliff’s δ of 0.98–1.00 at Days 10–14 (Table 2; Figures 3–5). Differences at Days 10, 12 and 14 remained significant after Holm adjustment (adjusted p≤0.001), whereas the Day 8 difference did not (adjusted p=0.087). Wound-level analyses and longitudinal models with cluster-robust standard errors also supported significant differences at Days 10–14 (all p<0.001). No significant differences between XWRAP and the pooled comparators were observed at Days 2, 4 or 6.

**Figure 3.**
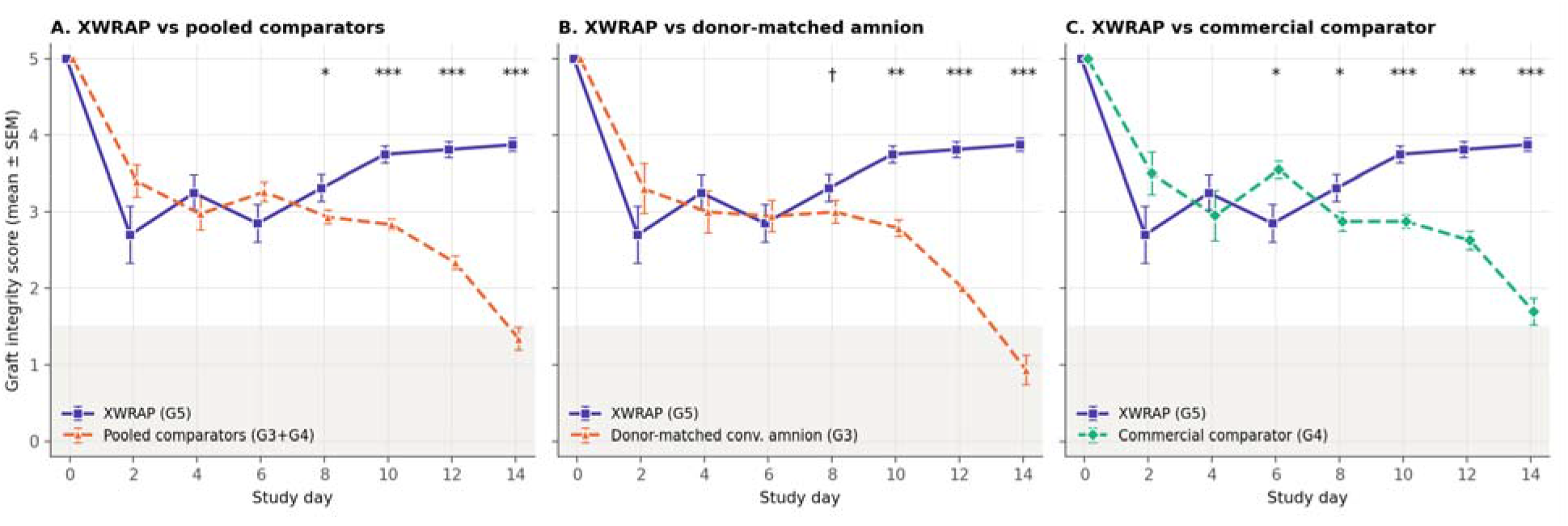
Graft-integrity trajectories of XWRAP compared with (A) the pooled comparator groups, (B) donor-matched conventional amnion, and (C) the commercial comparator (mean ± SEM). Symbols indicate animal-level Mann–Whitney comparisons at each visit: * p<0.05, ** p<0.01, *** p<0.001, † 0.05≤p<0.10.

**Figure 4.**
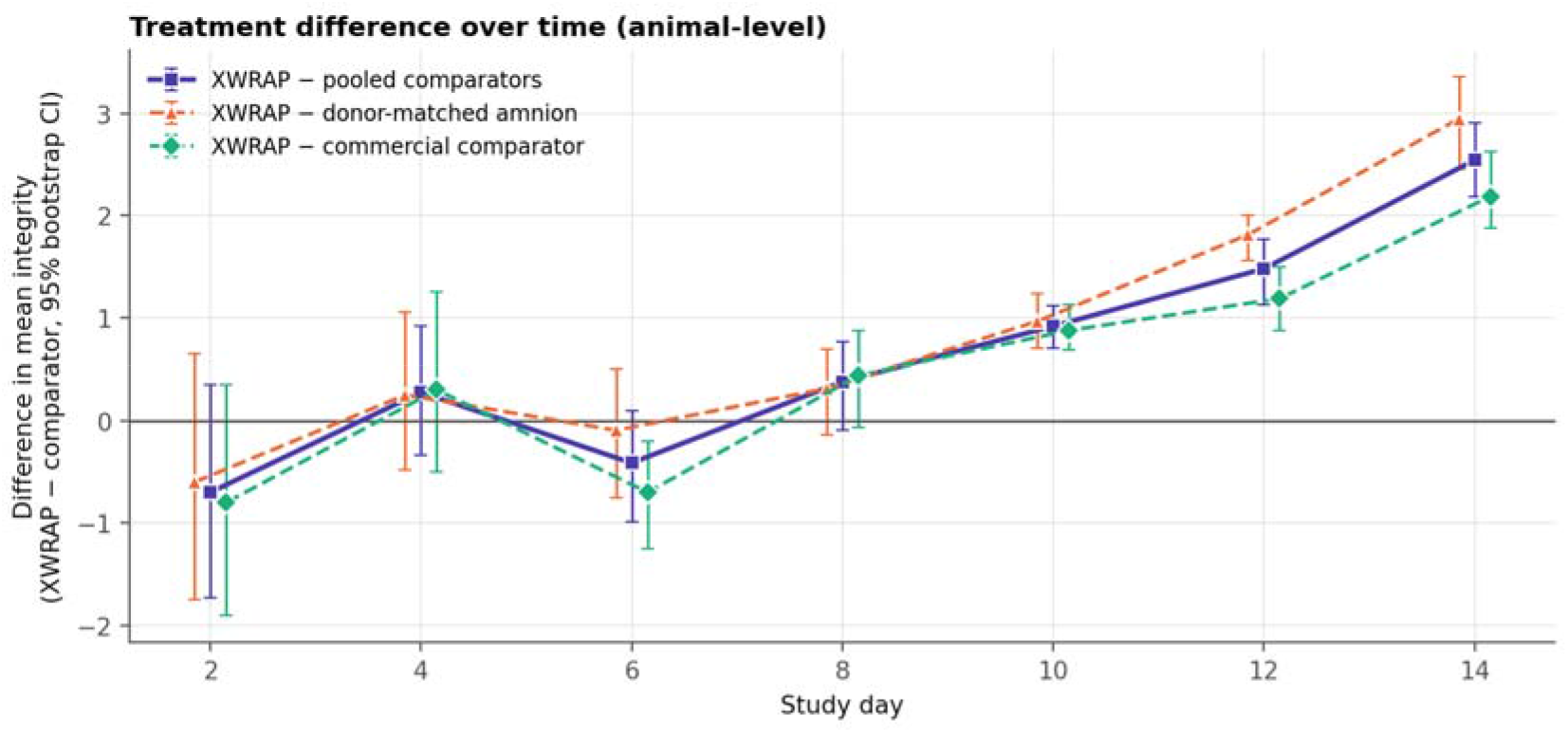
Difference in mean graft integrity between XWRAP and each comparator over time (animal-level means with 95% bootstrap confidence intervals). Values above zero indicate greater integrity with XWRAP.

**Figure 5.**
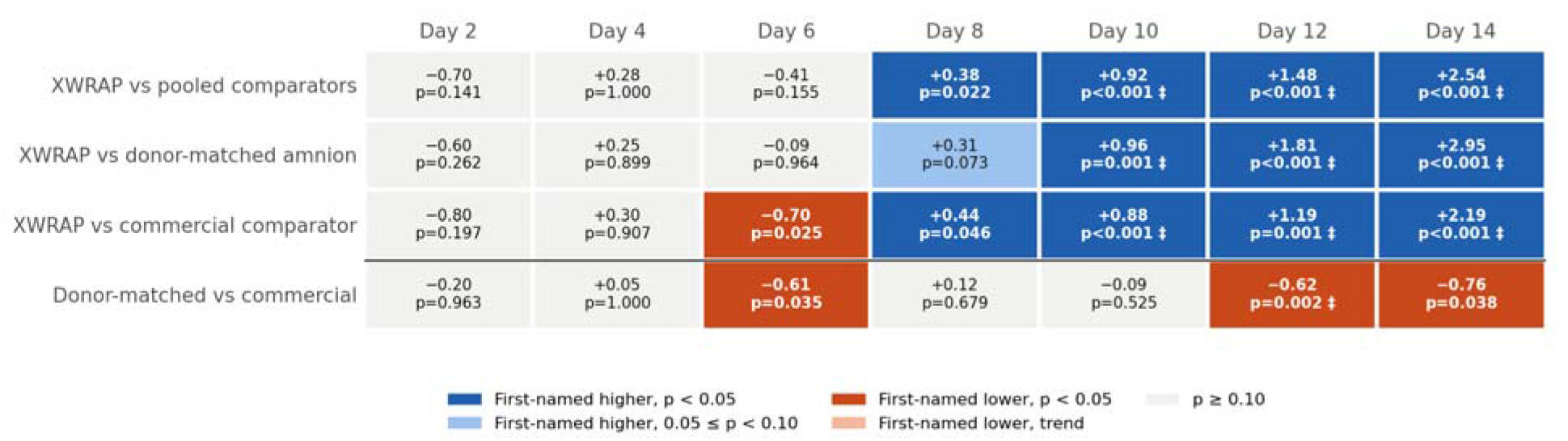
Summary of per-visit comparisons. Each cell shows the mean difference (first-named group minus second) and the animal-level Mann–Whitney p-value; dark shading indicates p<0.05, light shading 0.05≤p<0.10, and ‡ indicates significance after Holm adjustment across visits.

**Table 2.** Per-visit comparison of graft integrity between XWRAP and the comparator groups (animal-level means; two-sided Mann–Whitney U test). Holm-adjusted p-values across the seven visits are shown in parentheses for the individual comparators.

| Visit | XWRAP | Pooled comparators | Difference [95% CI] | Cliff's $\delta$ | p | Holm p | Donor-matched amnion | p donor-matched (Holm) | vs. Commercial comparator | p vs. commercial (Holm) |
| --- | --- | --- | --- | --- | --- | --- | --- | --- | --- | --- |
| Day 2 | 2.70 | 3.40 | –0.70 [–1.73, +0.35] | –0.30 | 0.141 | 0.424 | 3.30 | 0.262 (0.786) | 3.50 | 0.197 (0.394) |
| Day 4 | 3.25 | 2.97 | +0.28 [–0.34, +0.92] | 0.00 | 1.000 | 1.000 | 3.00 | 0.899 (1.000) | 2.95 | 0.907 (0.907) |
| Day 6 | 2.85 | 3.26 | –0.41 [–0.99, +0.09] | –0.31 | 0.155 | 0.424 | 2.94 | 0.964 (1.000) | 3.55 | 0.025 (0.098) |
| Day 8 | 3.31 | 2.93 | +0.38 [–0.10, +0.76] | 0.56 | 0.022 | 0.087 | 3.00 | 0.073 (0.291) | 2.88 | 0.046 (0.137) |
| Day 10 | 3.75 | 2.83 | +0.92 [+0.70, +1.12] | 1.00 | <0.001 | <0.001 | 2.79 | 0.001 (0.005) | 2.88 | <0.001 (0.004) |
| Day 12 | 3.81 | 2.33 | +1.48 [+1.13, +1.77] | 0.98 | <0.001 | <0.001 | 2.00 | <0.001 (0.004) | 2.63 | 0.001 (0.005) |
| Day 14 | 3.88 | 1.33 | +2.54 [+2.18, +2.90] | 1.00 | <0.001 | <0.001 | 0.93 | <0.001 (0.005) | 1.69 | <0.001 (0.004) |

From Day 10 through Day 14, XWRAP demonstrated substantial and consistent separation from both comparator products. Compared with donor-matched conventional amnion, XWRAP demonstrated significantly greater graft integrity at Day 10 (3.75 vs. 2.79; p=0.001), Day 12 (3.81 vs. 2.00; p<0.001), and Day 14 (3.88 vs. 0.93; p<0.001). At Day 8, XWRAP showed a trend toward greater integrity than donor-matched conventional amnion (3.31 vs. 3.00; p=0.073).

Compared with the commercial comparator, XWRAP demonstrated greater graft integrity at Day 8 (3.31 vs. 2.88; p=0.046), Day 10 (3.75 vs. 2.88; p<0.001), Day 12 (3.81 vs. 2.63; p=0.001), and Day 14 (3.88 vs. 1.69; p<0.001). The magnitude of these late-phase differences was substantial. Cliff’s δ ranged from 0.95 to 1.00 during the principal Day 10–14 comparisons, indicating near-complete separation between XWRAP-treated animals and comparator-treated animals. For both individual comparators, differences at Days 10–14 remained significant after Holm adjustment (adjusted p≤0.006), whereas the Day 8 differences did not.

Across predefined intervals, mean XWRAP integrity exceeded that of the pooled comparator groups during Days 8–10 (3.53 vs. 2.88; p<0.001), Days 10–14 (3.81 vs. 2.17; p<0.001), and the full Day 2–14 interval (3.46 vs. 2.75; p<0.001), but not during Days 2–6 (2.93 vs. 3.20; p=0.227). Between Day 6 and Day 14, mean XWRAP integrity changed by +0.88 points, compared with −2.03 points for the pooled comparator groups (p<0.001) (Table 3).

**Table 3.** Graft-persistence endpoints by treatment group (animal-level mean ± SD or mean; animal followed to Day 14 for slopes and Day 14 values) and Mann–Whitney comparisons with XWRAP.

| Endpoint | XWRAP | Donor-matched amnion | Commercial comparator | Pooled comparators | p vs. pooled | p vs. donor-matched | p vs. commercial |
| --- | --- | --- | --- | --- | --- | --- | --- |
| Original-graft degradation slope (units/day) | -0.036 ± 0.026 | -0.250 ± 0.035 | -0.187 ± 0.032 | -0.213 ± 0.045 | <0.001 | 0.003 | <0.001 |
| Graft integrity at Day 14 | 3.88 ± 0.23 | 0.93 ± 0.61 | 1.69 ± 0.53 | 1.33 ± 0.67 | <0.001 | <0.001 | <0.001 |
| Original-graft integrity at Day 14 (wound mean) | 3.92 | 0.90 | 1.83 | 1.41 | <0.001 | 0.003 | 0.001 |
| Mean integrity, Days 2–6 | 2.93 | 3.06 | 3.33 | 3.20 | 0.227 | 0.650 | 0.121 |
| Mean integrity, Days 8–10 | 3.53 | 2.89 | 2.88 | 2.88 | <0.001 | 0.003 | 0.002 |
| Mean integrity, Days 10–14 | 3.81 | 1.90 | 2.40 | 2.17 | <0.001 | 0.001 | <0.001 |
| Mean integrity, Days 2–14 | 3.46 | 2.65 | 2.83 | 2.75 | <0.001 | 0.002 | 0.006 |
| Change in integrity, Day 6 to Day 14 | 0.88 | -2.29 | -1.81 | -2.03 | <0.001 | 0.001 | <0.001 |

Between the two comparator groups, the commercial comparator retained greater integrity than donor-matched conventional amnion at Day 6 (3.55 vs. 2.94; p=0.035), Day 12 (2.63 vs. 2.00; p=0.002) and Day 14 (1.69 vs. 0.93; p=0.038).

### Original-Graft Analysis Confirmed Greater XWRAP Persistence

Because placement of a replacement graft could influence subsequent integrity scores, an original-graft analysis was performed in which each wound was censored following its first reapplication. The original-graft analysis reproduced the same overall pattern. At Day 14, mean original-graft integrity was: XWRAP: 3.92; Donor-matched conventional amnion: 0.90; and Commercial comparator: 1.83. Longitudinal degradation modeling similarly demonstrated a substantial difference in XWRAP’s degradation trajectory. The treatment-by-day interaction associated with the XWRAP group was +0.233 integrity-score units/day relative to the comparator groups (p<0.001). Thus, the observed persistence difference remained when replacement grafts were excluded from the analysis.

In per-visit original-graft comparisons, XWRAP integrity exceeded that of the pooled comparator groups at Days 10, 12 and 14 (all p<0.001), but not at Day 8 (p=0.192). The 95% confidence interval for the treatment-by-day interaction was 0.143 to 0.323 units/day. Mean original-graft degradation slopes were −0.036 ± 0.026 integrity units/day for XWRAP, compared with −0.213 ± 0.045 for the pooled comparator groups (−0.250 ± 0.035 for donor-matched conventional amnion and −0.187 ± 0.032 for the commercial comparator; p<0.001 vs. pooled, p=0.003 vs. donor-matched, p<0.001 vs. commercial) (Table 3; Figure 6). Time to first original-graft reapplication (score ≤1) did not differ significantly between XWRAP and the pooled comparators (log-rank p=0.14); all six XWRAP original-graft reapplications occurred at Days 2–4, whereas comparator failures were divided between Days 2–4 (n=10) and the terminal Day 14 assessment (n=11).

**Figure 6.**
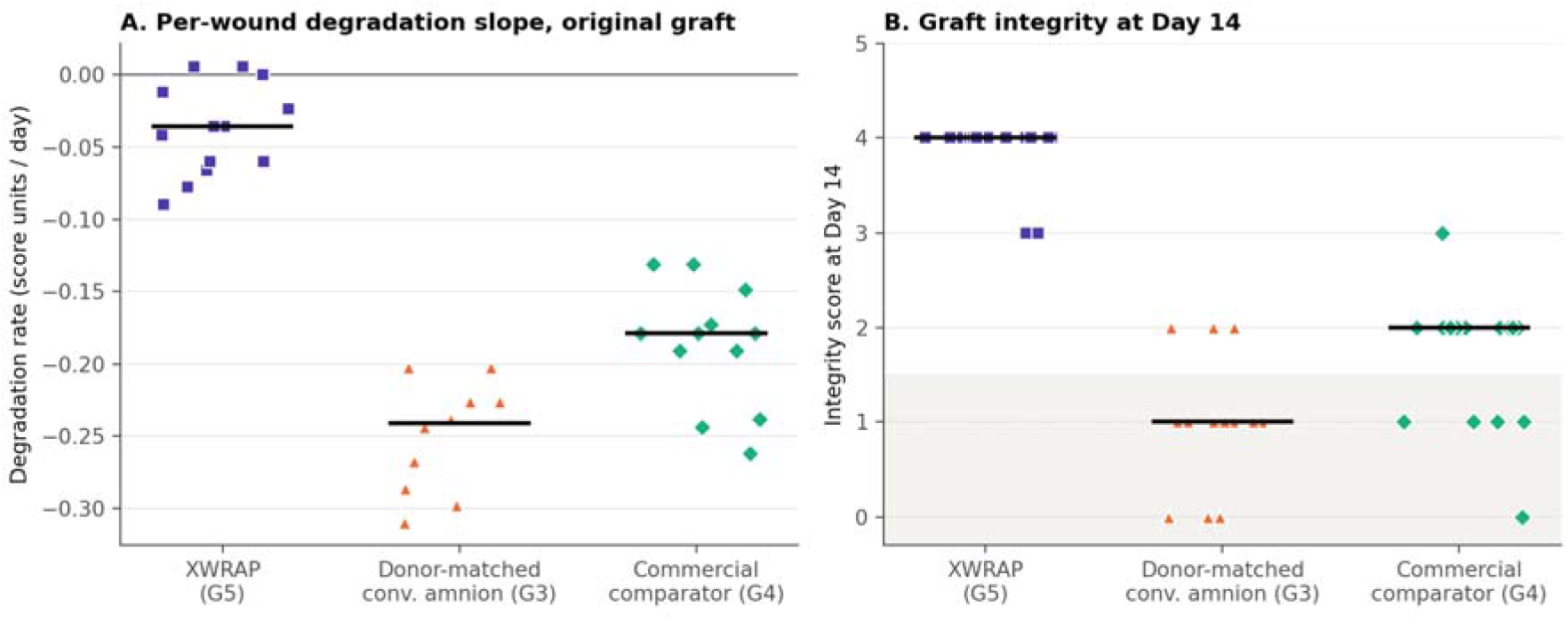
Per-wound original-graft degradation slopes (A) and graft-integrity scores at Day 14 (B) in wounds followed to Day 14. Horizontal bars indicate medians.

### Day 14 Reapplication Threshold

The practical significance of the persistence difference became particularly apparent at the terminal Day 14 assessment. The protocol-defined threshold for reapplication was an integrity score ≤1. At Day 14: XWRAP was 0/16 wounds (0%); Commercial comparator was 5/16 wounds (31%); and Donor-matched conventional amnion was 11/14 wounds (79%). The proportion of wounds at or below the threshold was significantly lower with XWRAP than with the pooled comparator groups (0/16 vs. 16/30 [53%]; Fisher’s exact p<0.001), donor-matched conventional amnion (0/16 vs. 11/14; p<0.001), and the commercial comparator (0/16 vs. 5/16; p=0.043).

Thus, 100% of XWRAP-treated wounds remained above the protocol-defined reapplication threshold at Day 14, compared with 69% of wounds treated with the commercial comparator and only 21% of wounds treated with donor-matched conventional amnion. Because Day 14 represented the terminal study visit, wounds meeting the threshold at this timepoint could not receive an additional graft. Consequently, the study design prevented this late-phase divergence from being reflected in the observed reapplication count. Had treatment continued under the same protocol-defined criterion, substantially more comparator-treated wounds would have been eligible for another graft at Day 14, while no XWRAP-treated wound had reached that threshold.

### Documented Reapplications and Dressing-Associated Graft Loss

Across the broader study, all documented reapplications occurred during Days 2–6. No graft reapplications occurred after Day 6. Seven reapplications occurred in the XWRAP group. Five of these seven events were documented in association with adhesion of the graft to the gauze dressing. The distinction between dressing-associated graft loss and in-situ biodegradation is relevant when considering mechanism. From an application-utilization perspective, however, either mechanism can result in loss of adequate graft coverage and the requirement for another graft.

In the comparator groups, 8 reapplications occurred with donor-matched conventional amnion (3 associated with dressing adhesion) and 6 with the commercial comparator (4 associated with dressing adhesion). Reapplication frequency was therefore similar for XWRAP and the pooled comparator groups (3.24 vs. 3.40 per 100 wound-days; rate ratio 0.95, 95% CI 0.33–2.53; p=1.00; cluster-bootstrap 95% CI 0.21–3.25), and did not differ significantly from donor-matched conventional amnion (rate ratio 0.79; p=0.80) or the commercial comparator (rate ratio 1.17; p=1.00). The proportion of wounds receiving at least one reapplication was 25% in both XWRAP-treated wounds (5/20) and pooled comparator wounds (10/40). No significant difference was observed at any individual visit (all p≥0.14), and excluding dressing-associated events did not change this result (rate ratio 0.54; p=0.73) (Table 4; Figure 7).

**Figure 7.**
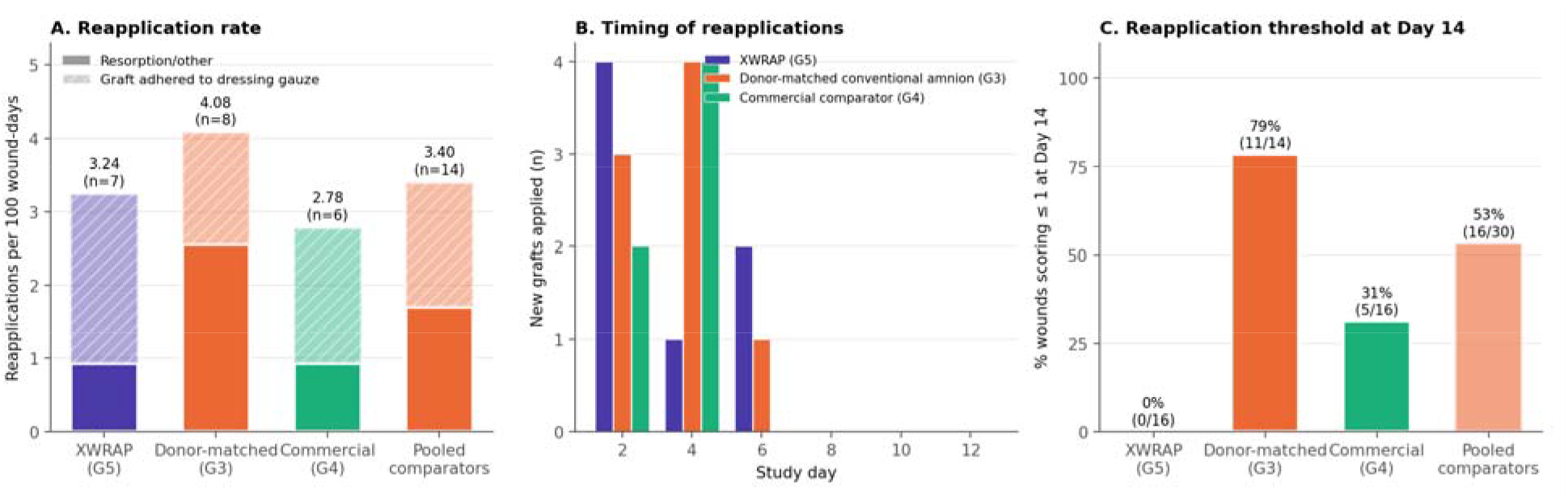
(A) Documented reapplication rate per 100 wound-days, divided into dressing-associated (hatched) and other events. (B) Timing of reapplications. (C) Proportion of wounds at or below the reapplication threshold (score ≤1) at the terminal Day 14 assessment.

**Table 4.** Documented graft reapplications (Days 2–12) and wounds at or below the reapplication threshold at Day 14. XWRAP vs. pooled comparators: reapplication rate ratio 0.95 (95% CI 0.33–2.53), p=1.00; Day 14 threshold, Fisher’s exact p<0.001.

| Group | Wounds | Wound-days at risk | Reapplications (n) | Dressing-associated (n) | Per 100 wound-days | Wounds with $\geq 1$ reapplication | Wounds $\leq 1$ at Day 14 |
| --- | --- | --- | --- | --- | --- | --- | --- |
| XWRAP (G5) | 20 | 216 | 7 | 5 | 3.24 | 5 (25%) | 0/16 (0%) |
| Donor-matched conventional amnion (G3) | 20 | 196 | 8 | 3 | 4.08 | 6 (30%) | 11/14 (79%) |
| Commercial comparator (G4) | 20 | 216 | 6 | 4 | 2.78 | 4 (20%) | 5/16 (31%) |
| Pooled comparators (G3+G4) | 40 | 412 | 14 | 7 | 3.40 | 10 (25%) | 16/30 (53%) |

The timing of these events is particularly important. Early reapplications occurred before the substantial late-phase divergence in graft persistence. The largest separation between XWRAP and the comparator groups emerged during Days 10–14, after the final study visit at which reapplication occurred. Consequently, the observed reapplication counts during the study do not capture the potential utilization consequence of the pronounced late-phase persistence difference.

## Discussion

### Proprietary Processing Produced a Distinct Graft-Persistence Profile

The principal finding of the present analysis is that XWRAP demonstrated markedly greater late-phase persistence than either donor-matched conventionally processed amnion or a commercial placental tissue comparator.

The donor-matched comparison is particularly informative. XWRAP and the conventional comparator originated from the same amnion donor, yet their persistence trajectories diverged substantially during the later study period. At Day 14, mean graft-integrity scores were 3.88 and 0.93, respectively. This observation supports the concept that tissue source alone does not determine product performance. Manufacturing and processing methodology can materially influence the physical behavior of human amniotic membrane following application. [18–30, 32]

The commercial comparator provided an independent reference. At Day 14, its mean integrity score was 1.69—greater than donor-matched conventional amnion but substantially lower than XWRAP. The consistency of these findings across donor-matched and commercial comparisons supports a distinct late-phase persistence phenotype associated with the proprietary processed allograft.

### Molecular Basis for ECM Stabilization and Sustained Graft Persistence

The substantial differences in graft persistence observed in this study are biologically plausible given the molecular architecture of the extracellular matrix and the established effects of tissue processing on collagen stability.

Collagen is organized hierarchically. Individual polypeptide chains assemble into triple-helical collagen molecules, which subsequently associate into fibrils and higher-order extracellular matrix structures. [21, 22] The stability of this network depends in part upon intermolecular interactions and naturally occurring covalent crosslinks between adjacent collagen molecules. These molecular connections contribute to the mechanical strength, structural organization, and resistance to enzymatic degradation characteristic of mature collagen-rich tissues. [21–25]

Chemical modification provides a means of altering this molecular architecture. Multiple crosslinking approaches have been described in the biomaterials literature, including glutaraldehyde, carbodiimide-based chemistries, genipin, and naturally occurring polyphenols. Although the reaction mechanisms differ, these approaches can increase intermolecular stabilization within collagen-rich matrices by promoting additional covalent interactions between reactive sites on adjacent proteins. [23–30]

At the molecular level, increased intermolecular stabilization can reduce the mobility and separation of collagen molecules within the matrix and increase the energetic and enzymatic requirements for matrix disruption. Proteolytic enzymes must not only recognize and cleave susceptible peptide sequences but operate within a more highly interconnected molecular network. Consequently, modification of intermolecular collagen architecture can alter susceptibility to proteolytic cleavage and slow the progressive loss of matrix structural integrity. [21, 24–30]

This characteristic may be particularly relevant within wound environments. Matrix metalloproteinases and other proteolytic enzymes participate normally in extracellular matrix turnover during tissue repair. In chronic wounds, however, elevated proteolytic activity can accelerate degradation of extracellular matrix proteins and matrix-associated signaling molecules. [2,4,6,7] A matrix that loses structural integrity prematurely may therefore provide physical coverage and matrix-associated biological interactions for a shorter interval.

The objective of extracellular matrix stabilization is not simply to maximize resistance to degradation. Excessive crosslink density or inappropriate chemical modification can increase matrix rigidity, decrease porosity, alter cellular recognition sites, interfere with diffusion or release of matrix-associated molecules, and impede cellular infiltration and host-mediated remodeling. The biologically desirable state is therefore a balance in which the matrix remains sufficiently stable during active tissue repair while retaining the capacity to interact with cells and undergo progressive remodeling. [26–30]

The findings of the present study are consistent with such a balance. XWRAP demonstrated substantially greater late-phase graft persistence than both donor-matched conventionally processed amnion and a commercial placental tissue comparator. At Day 14, mean XWRAP graft-integrity score was 3.88 compared with 0.93 for donor-matched conventional amnion, representing more than a four-fold difference in mean integrity score.

Importantly, the increased persistence observed in the present analysis accompanies the favorable wound-healing biology demonstrated in our previously published study [1]. XWRAP was associated with increased proliferative activity, reduced tissue injury, favorable inflammatory remodeling, and more advanced progression toward wound resolution relative to conventionally processed placental tissue.

Taken together, these observations support a model in which proprietary processing alters extracellular matrix characteristics sufficiently to prolong structural persistence while preserving the biological interactions associated with tissue repair. Further controlled studies of proteolytic degradation, collagen architecture, and matrix-associated protein retention will help define the molecular mechanisms underlying this phenotype.

### Greater Persistence Was Accompanied by Favorable Wound-Healing Biology

The significance of the persistence findings becomes more apparent when considered together with our previously published analysis of wound-healing biology in this model [1]. In that study, XWRAP demonstrated a more favorable wound-healing phenotype than conventionally processed placental tissue across multiple independent measures of tissue repair. Histologic and immunohistochemical evaluation demonstrated increased proliferative activity, reduced ulceration severity, reduced giant-cell response, favorable inflammatory remodeling, and more advanced progression toward wound resolution [1]. These findings are important because they demonstrate that the XWRAP persistence phenotype exists alongside favorable biological performance. Rather than simply remaining physically present for a longer period, XWRAP demonstrated both sustained graft integrity and a more favorable wound-healing biologic profile.

This combination provides a broader framework for evaluating placental tissue performance. A graft that degrades rapidly may provide biologically active tissue for only a limited interval. Conversely, persistence without favorable biological activity would provide limited evidence of regenerative value. An optimized placental tissue allograft would ideally combine sufficient matrix persistence with favorable cellular, inflammatory, and tissue-remodeling responses. The combined findings from these analyses suggest that proprietary processing can influence both dimensions.

### Graft Persistence Provides a Mechanistic Basis for Reduced Application Requirements

Application frequency is an important practical characteristic of placental tissue products. When a graft is no longer adequately present at the wound interface, another application may be required to restore coverage. The number of applications required during a course of wound management may therefore be influenced not only by the biological activity of the product, but also by the duration for which each application remains sufficiently intact. The present study provides direct experimental evidence of substantial differences in this characteristic.

At Day 14, 100% of XWRAP-treated wounds remained above the protocol-defined reapplication threshold. In contrast, only 21% of wounds treated with donor-matched conventional amnion and 69% of wounds treated with the commercial comparator remained above that threshold. Stated differently, 79% of donor-matched conventional wounds and 31% of commercial-comparator wounds had reached the point at which another application would have been indicated under the study protocol, compared with 0% of XWRAP-treated wounds.

The study ended at precisely the point at which these differences were greatest. Because Day 14 was terminal, the comparator wounds meeting the reapplication criterion could not receive another graft. Consequently, the observed number of applications during the study necessarily underestimates the potential application-utilization consequences of the late-phase persistence differences. The data therefore do not establish the number of applications that would be required to achieve wound closure in human chronic wounds. They do, however, establish a clear experimental basis for the hypothesis that a graft capable of remaining intact longer may require fewer repeat applications over a course of wound management. [35]

### Persistence and Wound-Healing Biology May Act Together to Influence Application Efficiency

Graft persistence and wound-healing biology should not necessarily be considered independent characteristics. The duration for which an extracellular matrix remains present determines the period during which it can provide physical coverage and interact with the local tissue environment. At the same time, the biological response occurring during that interval influences progression of the underlying wound.

A product capable of maintaining graft integrity for a longer period while simultaneously supporting a favorable wound-healing trajectory could therefore have an important practical advantage: each application may remain present through a greater portion of the repair process while the underlying tissue progresses toward resolution.

This provides a mechanistic framework through which sustained graft persistence and favorable wound-healing biology may act together to support faster healing with fewer applications. Our previously published findings demonstrated a more favorable wound-healing biologic profile with XWRAP relative to conventionally processed placental tissue [1]. The present analysis demonstrates the complementary finding that XWRAP also persists substantially longer than donor-matched conventional amnion and a commercial placental tissue comparator.

Taken together, these findings support a product-performance profile combining favorable wound-healing biology with sustained graft persistence. A graft that remains present through a greater portion of the repair interval while supporting a more favorable healing trajectory provides a mechanistic and experimental rationale for the hypothesis that faster wound healing may be achieved with fewer applications.

This hypothesis requires confirmation in prospective clinical studies measuring application number, application interval, time to closure, and total applications required to achieve closure. Nevertheless, the combined experimental findings establish the two components underlying this hypothesis: favorable wound-healing biology and sustained graft persistence.

### Economic Implications of Faster Healing With Fewer Applications

The economic value of an advanced wound-care product cannot be determined solely by its acquisition price. The more relevant measure is the total cost required to achieve durable wound closure. This includes the cost of the product, resources associated with each application, the duration of ongoing wound management, and the potential downstream costs associated with an ulcer that remains open.

This distinction is particularly relevant for diabetic foot ulcers. Published U.S. Medicare analyses demonstrate that beneficiaries with diabetic foot ulcers utilize substantially greater healthcare resources than matched beneficiaries with diabetes but without foot ulcers, including 138% more hospital days, 85% more home-health days, 41% more emergency-department visits, and 35% more outpatient and physician visits. Incremental annual Medicare expenditures associated with diabetic foot ulcers have been estimated at approximately $11,710 per patient. [33]

Other U.S. analyses demonstrate that the economic burden increases substantially as wounds become more severe and remain open longer. In a multidisciplinary diabetic limb-preservation program, mean total cost was approximately $24,226 per wound episode and mean time to healing was approximately 136 days, with costs increasing markedly with wound severity. [34]

These observations suggest that a product capable of supporting faster wound resolution with fewer applications may influence cost through three complementary mechanisms: (1) Fewer application encounters, reducing product acquisition and application-associated resource utilization; (2) Shorter duration of open-wound management, reducing ongoing wound-care resources until closure; and (3) Reduced exposure to downstream complications, including infection, hospitalization, surgical intervention, and amputation. [35]

The present animal studies were not designed to establish clinical cost savings. The following analyses are therefore presented as illustrative economic scenarios intended to demonstrate the potential magnitude of these effects and identify parameters for prospective clinical and economic evaluation.

### Application and Routine Wound-Care Scenario

For purposes of scenario modeling, a graft acquisition cost of $500 per application was assumed. An additional $500 per application encounter was assumed to represent wound assessment, wound-bed preparation and debridement when indicated, provider and facility resources, dressings, supplies, and related clinical services. The modeled total cost of each graft-application encounter was therefore: $500 graft + $500 application-associated resources = $1,000 per application encounter. A separate cost of $300 per non-application open-wound week was assumed to represent ongoing wound-management resources during weeks in which the ulcer remains open but another graft is not applied, including dressing changes, supplies, wound assessment, off-loading management, and other outpatient or home-based wound care. These values are illustrative assumptions rather than measured costs from the present study or specific Medicare reimbursement rates.

Consider a conventional treatment scenario requiring four graft applications over eight weeks compared with a more efficient scenario requiring two applications with closure at four weeks. Conventional Treatment Scenario: 4 graft-application encounters × $1,000 = $4,000; 4 additional open-wound weeks × $300 = $1,200; and Total routine wound-management cost = $5,200. Faster-Healing/Fewer-Application Scenario: 2 graft-application encounters × $1,000 = $2,000; 2 additional open-wound weeks × $300 = $600; and Total routine wound-management cost = $2,600. Under these illustrative assumptions, reducing a treatment course from four applications over eight weeks to two applications with closure at four weeks would reduce modeled routine episode-of-care cost from $5,200 to $2,600, a 50% reduction.

Importantly, graft acquisition costs are included in both treatment pathways. The modeled savings therefore reflect the total cost of achieving wound closure—including the cost of the grafts themselves, application-associated resources, and ongoing wound management—and illustrate how faster healing with fewer applications may reduce overall episode-of-care costs even after accounting for product acquisition costs.

### Sensitivity to Graft Acquisition Cost

Because product acquisition price can vary substantially, the economic model was additionally evaluated across a range of hypothetical graft prices while holding the other assumptions constant. Under the two-application/four-week scenario, modeled routine episode cost would be: $2,600 at $500 per graft; $3,100 at $750 per graft; $3,600 at $1,000 per graft; $4,100 at $1,250 per graft; $4,600 at $1,500 per graft; and $5,200 at $1,800 per graft. Under these assumptions, a two-application treatment pathway remains less costly than the $5,200 conventional reference pathway even when the acquisition cost of each graft is substantially greater. This illustrates an important economic principle: a higher-performing graft may generate lower total episode-of-care costs even at a higher unit acquisition price if fewer applications and a shorter duration of wound management are required. [35]

### Potential Economic Impact of Downstream Complications

Routine wound-management costs represent only one component of the economic burden associated with diabetic foot ulcers. The more consequential costs frequently arise when an open ulcer progresses to infection, hospitalization, surgical intervention, or lower-extremity amputation. Published U.S. hospital analyses have reported mean costs of approximately $11,290 for hospitalization involving an infected diabetic foot ulcer, with inpatient costs increasing to approximately $14,270 for minor amputation and $18,877 for major amputation. [38] Medicare claims analyses have additionally reported lower-extremity amputation in approximately 3.8% of beneficiaries with diabetic foot ulcers over a 12-month period. [33]

Earlier durable closure may therefore create economic value beyond reducing routine wound-care encounters because it shortens the period during which the patient remains exposed to complications arising from an open ulcer. The present experimental studies do not establish the magnitude by which XWRAP would reduce infection, hospitalization, or amputation risk in humans. These potential benefits are therefore appropriately evaluated through sensitivity analysis.

Using the published 3.8% annual amputation incidence and published index-hospitalization costs, the expected hospitalization burden associated with amputation is approximately: 3.8% × $14,270 = $542 per DFU patient at the lower end, and 3.8% × $18,877 = $717 per DFU patient at the upper end. If earlier durable closure hypothetically reduced this risk exposure by 25%, the expected savings from index amputation hospitalization alone would be approximately $136–$179 per patient. At a hypothetical 50% reduction, expected savings would be approximately $271–$359 per patient. These estimates exclude rehabilitation, prosthetic care, recurrent hospitalization, home-health utilization, subsequent procedures, long-term disability, and other costs following amputation and are therefore intentionally conservative. Infection-related hospitalization represents an additional major economic burden. Because infection, hospitalization, and amputation can occur within the same clinical episode, these costs should not simply be added as independent events; doing so could double count the same complication pathway. Accordingly, complication avoidance is most appropriately considered as a probability-weighted sensitivity analysis layered onto routine episode-of-care costs.

### Integrated Per-Patient Economic Scenario

Under the illustrative routine-care model the conventional treatment pathway costs $5,200 and the faster-healing/fewer-application pathway costs $2,600 with a routine-care savings of $2,600 per wound episode, or 50%. Adding the conservative hypothetical 25% reduction in amputation-risk exposure increases modeled net savings to approximately $2,736–$2,779 per wound episode. At a hypothetical 50% reduction in amputation-risk exposure, modeled savings increase to approximately $2,871–$2,959 per wound episode. These calculations still exclude potential savings associated with avoided infection-related hospitalization and the substantial post-amputation costs of rehabilitation, prosthetic care, home health, recurrent hospitalization, and disability. The modeled savings should therefore not be interpreted as predicted XWRAP outcomes. Rather, they demonstrate how faster durable closure can generate economic value through multiple pathways.

### Potential Health-System Impact

The economic implications become increasingly meaningful when applied across a hospital system or integrated delivery network. Using only the conservative $2,600 routine-care savings per wound episode and excluding downstream complication savings with 100 eligible wound episodes annually there is potential savings of $260,000, $1.3 million potential savings with 500 eligible wound episodes, $2.6 million potential savings with 1,000 eligible wound episodes and $13.0 million potential savings with 5,000 eligible wound episodes.

Using the integrated scenario incorporating the illustrative downstream-event reduction would increase these amounts further. For example, at approximately $2,750–$2,950 in modeled net savings per episode, a health system treating 1,000 eligible wounds annually would generate approximately $2.75–$2.95 million in modeled annual savings. These calculations illustrate why relatively modest improvements in healing and application efficiency can become economically significant across large health systems and group purchasing organizations.

### Potential Medicare Budget Impact

At the national level, the opportunity is substantially larger. CMS has estimated that diabetic foot ulcers account for approximately $6.2 billion to $18.7 billion in annual Medicare expenditures, depending upon the methodology used to attribute wound-related healthcare utilization. [36] Rather than assuming that the illustrative per-patient model applies universally, a conservative national sensitivity analysis can evaluate the potential impact of relatively small net reductions in the established Medicare DFU burden. A 5% reduction in total Medicare DFU expenditures would generate approximately $310–$935 million in annual savings. Savings would increase to approximately $620 million–$1.87 billion at 10%, $930 million–$2.81 billion at 15%, $1.24–$3.74 billion at 20%, and $1.55–$4.68 billion at 25%. These estimates represent net reductions in total DFU expenditures, inclusive of the cost of the treatment strategy responsible for improved outcomes; they do not assume that the therapy itself is cost-free.

A complementary bottom-up calculation provides an additional perspective. If a more efficient wound-management strategy produced only $1,000 in net savings per treated episode after accounting for therapy acquisition cost, every 100,000 treated episodes would correspond to approximately $100 million in healthcare savings. At the illustrative $2,600 net routine-care savings per episode, 100,000 treated episodes would correspond to approximately $260 million in modeled savings, 500,000 episodes to approximately $1.3 billion, and 1 million episodes to approximately $2.6 billion. These bottom-up estimates are directionally consistent with the percentage-based sensitivity analysis applied to the published Medicare DFU expenditure burden.

### Implications for Value-Based Wound Care

The economic implications extend beyond the acquisition price of an individual placental tissue allograft. The relevant economic endpoint for a health system or payer is ultimately: What does it cost to achieve durable wound closure? A lower-priced product that requires repeated applications or is associated with prolonged wound management may generate greater total episode-of-care expenditures than a product with a higher acquisition price that achieves faster closure with fewer applications. [34,35] Conversely, a product capable of supporting favorable wound-healing biology while remaining present through a greater portion of the repair interval could potentially reduce product utilization, treatment encounters, duration of wound management, and exposure to costly downstream complications.

The experimental findings presented here provide a rationale for evaluating this hypothesis clinically. XWRAP demonstrated more than four times the mean Day 14 graft-integrity score of donor-matched conventionally processed amnion, and no XWRAP-treated wound reached the protocol-defined reapplication threshold at Day 14 compared with 79% of donor-matched conventional wounds. Our previously published analysis demonstrated the complementary finding of a more favorable wound-healing biologic profile with XWRAP. Together, these findings support prospective evaluation of a clinically and economically important hypothesis: Can sustained graft persistence and favorable wound-healing biology produce faster durable closure with fewer applications and lower total episode-of-care cost? If confirmed in prospective clinical studies, the potential economic implications could extend from individual wound episodes to substantial savings across hospital systems, integrated delivery networks, and the Medicare program.

### Implications for Placental Tissue Product Differentiation

Placental tissue products are frequently grouped according to common tissue source. The present findings, together with our previously published wound-healing analysis, suggest that tissue source alone provides an incomplete characterization of product performance. XWRAP and the donor-matched conventional comparator originated from the same human amnion donor, yet demonstrated markedly different late-phase persistence profiles. By Day 14, XWRAP maintained a mean integrity score of 3.88 compared with 0.93 for the donor-matched conventional product. [9,12,16,18–20,32]

The previous biological analysis similarly demonstrated differences in wound-healing phenotype between proprietary processed and conventionally processed placental tissue. Together, these observations demonstrate that processing methodology can materially influence both the physical persistence and biological performance of placental tissue following implantation. [1,18–20,26,28–30,32] Accordingly, placental tissue allografts derived from the same tissue source should not necessarily be assumed to be functionally equivalent.

## Limitations

Several limitations should be considered when interpreting these findings. First, the study utilized an acute full-thickness excisional wound model in healthy rats. This model permits controlled comparison of graft behavior but does not reproduce the complex pathophysiology of human chronic wounds, including diabetes, vascular disease, neuropathy, persistent inflammation, elevated protease activity, and other factors that may influence both graft degradation and wound repair. Second, graft integrity was evaluated using a semi-quantitative ordinal scoring system. Although longitudinal analyses demonstrated large and consistent treatment differences, direct biochemical measurement of matrix degradation was not performed. Controlled ex-vivo degradation studies will be useful for quantifying proteolytic degradation kinetics independently of dressing interactions and other in-vivo variables. Third, dressing-associated graft loss occurred during the early study period. Five of seven XWRAP reapplications were associated with graft adhesion to gauze. These events complicate mechanistic interpretation of early graft loss but do not alter the application-utilization consequence of requiring another graft. Importantly, the original-graft analysis independently confirmed the late persistence findings. Fourth, the study terminated at Day 14. This represents an important limitation for the reapplication analysis because the greatest treatment separation occurred at the terminal assessment. At Day 14, 79% of donor-matched conventional wounds and 31% of commercial-comparator wounds had reached the protocol-defined reapplication threshold, compared with no XWRAP-treated wounds, but no subsequent applications were permitted. Longer-duration studies are therefore needed to directly quantify how these late persistence differences affect total application requirements. Fifth, the present study evaluated finished placental tissue allografts rather than individual manufacturing variables. Accordingly, although substantial differences in graft persistence were observed between XWRAP and the comparator products, the study was not designed to isolate the relative contribution of individual proprietary processing parameters to the observed persistence phenotype. The findings should therefore be interpreted as demonstrating differences in finished-product performance rather than establishing a specific manufacturing mechanism responsible for those differences. Finally, the study demonstrates experimental graft persistence and wound-healing biology rather than clinical application efficiency. Prospective human studies will be necessary to determine whether the observed combination of favorable wound-healing biology and sustained graft persistence translates into fewer applications required to achieve wound closure.

## Future Directions

The present findings identify several opportunities for further investigation. Controlled ex-vivo experiments should directly compare degradation kinetics of proprietary processed and conventionally processed placental tissue under defined proteolytic conditions. Such studies would permit quantitative assessment of matrix stability independent of dressing adhesion, mechanical displacement, and observer-based scoring. Future in-vivo studies should extend beyond Day 14 and permit continued reapplication according to predefined integrity criteria. This would allow the late-phase persistence differences observed in the present study to translate into directly measured differences in total application utilization. Most importantly, prospective clinical studies should evaluate wound-healing trajectory, time to closure, application interval, and total applications required to achieve closure within the same patients. These endpoints would directly test the hypothesis generated by the present experimental findings: that a placental tissue allograft combining favorable wound-healing biology with sustained graft persistence may achieve successful wound management with fewer applications.

## Conclusion

The present study demonstrates that XWRAP, a proprietary processed human amniotic membrane allograft, exhibits substantially greater late-phase graft persistence than both donor-matched conventionally processed human amnion and a commercial placental tissue comparator. At Day 14, XWRAP maintained a mean graft-integrity score of 3.88, compared with 0.93 for donor-matched conventional amnion and 1.69 for the commercial comparator. Importantly, 100% of XWRAP-treated wounds remained above the protocol-defined reapplication threshold, whereas 79% of donor-matched conventional wounds and 31% of commercial-comparator wounds had reached the threshold for another application. The persistence phenotype remained evident when analysis was restricted to original grafts, demonstrating that the result was not dependent upon replacement material. These persistence findings complement our previously published analysis demonstrating that XWRAP also produced a more favorable wound-healing biologic profile than conventionally processed placental tissue across multiple measures of cellular proliferation, inflammation, tissue injury, remodeling, and progression toward wound resolution [1]. Taken together, the two studies support a combined product-performance profile in which XWRAP demonstrates both sustained graft persistence and favorable wound-healing biology. A graft that remains present through a greater portion of the repair interval while supporting a more favorable healing trajectory provides a mechanistic and experimental rationale for the hypothesis that faster wound healing may be achieved with fewer applications. These findings further demonstrate that placental tissue products derived from similar or identical tissue sources should not necessarily be considered functionally equivalent. Proprietary processing can materially influence both how a placental tissue allograft interacts with the wound environment and how long that allograft remains present following application. Prospective clinical studies evaluating wound closure together with application frequency are warranted to determine whether the combination of favorable wound-healing biology and sustained graft persistence observed with XWRAP translates into fewer applications required to achieve closure in human chronic wounds.

